# *D*-plus: Beyond the *D*-statistic — Accurate Detection and Characterization of Introgression Despite Among-Lineage Rate Variation

**DOI:** 10.64898/2026.10.01.755911

**Authors:** Xiao-Xu Pang, Da-Yong Zhang

## Abstract

Ubiquitous in phylogenomic and population-genomic introgression work, the *D*-statistic has fallen victim to its own success. Its reliability is compromised by two major confounding factors: among-lineage substitution rate heterogeneity, which generates spurious signals of introgression, and unsampled “ghost” lineages, which mislead the characterization of introgression by erroneously attributing ghost ancestry to extant species. Here, we developed *D*-plus, a maximum-likelihood tool that explicitly accounts for both confounding factors to unpack the sources of significant *D*-statistic signals. Simulations demonstrate that *D*-plus effectively mitigates false introgression calls under among-lineage rate variation. Meanwhile, it retains high sensitivity for genuine introgression signals and correctly assigns detected signals to introgression involving ghost lineages, or solely between extant non-sister lineages, or a combination of the two. Our *D*-plus re-analysis of phylogenomic data from *Jaltomata* (Solanaceae) overturns prior interpretations of significant *D*-statistic signals, which were conventionally regarded as evidence for recurrent non-sister introgression. Instead, our results favor alternative explanations invoking artefacts driven by lineage-specific rate heterogeneity as well as two ghost introgression events. Collectively, *D*-plus addresses major methodological limitations inherent to traditional *D*-statistic analyses and enables accurate detection and characterization of introgression events.

## Introduction

With genome-wide data accumulating for a growing array of species across the tree of life, unprecedented opportunities have emerged to investigate the prevalence and dynamics of interspecific introgression (Schumer and Rieseberg 2026). Among the tools developed for this purpose, Patterson’s *D*-statistic (Green et al. 2010) has emerged as a workhorse of phylogenomic and population-genomic studies because it provides a simple, computationally tractable test that can be applied to genome-scale data. The *D*-statistic quantifies counts of biallelic site patterns from a quartet of taxa with the topology (((P1, P2), P3), O). Taking the outgroup-observed allele as the ancestral state “*A*” and the alternative allele as the derived state “*B*”, two discordant site patterns — *ABBA* and *BABA* — are informative for this test. Under the null model of no gene flow, *ABBA* and *BABA* polymorphisms are thought to arise exclusively from incomplete lineage sorting (ILS) and are thus expected to occur at equal frequencies. Conversely, significant departures from this parity are conventionally interpreted as evidence of introgression between P3 and one of its non-sister lineages (P1 or P2). However, the full reasoning underlying *D*-statistic analyses implicitly relies on two idealized assumptions that frequently fail in empirical data, leading to systematic bias.

The first critical assumption is a strict molecular clock with uniform substitution rates across lineages. Molecular evolutionary rates vary substantially among lineages as a consequence of biological factors including generation time, effective population size, metabolic rate, and longevity (Bromham 2009; Thomas et al. 2010; Lanfear et al. 2014). In genomic datasets, technical processes can further generate lineage-specific distortions resembling rate variation: differences in genetic distance from the reference and uneven resequencing depth can produce lineage-specific mapping and genotyping errors (Akopyan et al. 2025; Bustos-Caparros et al. 2025; Ji et al. 2025; Maurstad et al. 2025). Such biological or artefactual rate heterogeneity can impair *D*-statistic inference: homoplasy (i.e., independent origins of identical alleles across lineages) provides an alternative source of discordant site patterns to ILS, through which rate variation can perturb the *ABBA-BABA* balance even without introgression. Recent theoretical and simulation studies have demonstrated that the *D*-statistic can be remarkably sensitive to this effect, with even modest rate differences between sister lineages producing significant signals in the absence of gene flow (Frankel and Ané 2023; Koppetsch et al. 2024; Pang et al. 2025). Koppetsch et al. (2024) address this challenge by exploiting the spatial distribution of *D*-informative sites to disentangle rate-variation artefacts from genuine introgression; nevertheless, their approach remains vulnerable to mapping bias and mutation hotspots.

The second foundational assumption is that introgression occurs solely among sampled extant lineages. Empirical analyses, however, are bounded by practical constraints, such as the unavailability of extinct or unknown species, and study-specific sampling priorities. This leaves a substantial share of unsampled “ghost” lineages capable of introducing genetic material into focal lineages through historical hybridization — a phenomenon known as ghost introgression (Ottenburghs 2020; Tricou et al. 2022) that is increasingly documented across taxa (Pawar et al. 2023; Zhang et al. 2024; Zhu et al. 2025; Gedman et al. 2026; Stanton et al. 2026; Yang et al. 2026; Zhang et al. 2026). Theoretical and simulation studies (Tricou et al. 2022; Pang and Zhang 2024) have shown that ghost introgression can produce *ABBA-BABA* imbalance, frequently causing misinterpretation of *D*-statistic results by misassigning ghost ancestry to extant species. Recently, Yang et al. (2026) and (Cheng et al. 2026) applied a Bayesian test in the BPP framework (Yang 2015; Flouri et al. 2020; Flouri et al. 2023) to separate ghost introgression and non-sister introgression, but the MCMC-based method is operationally complex and computationally expensive, restricting its use in large-scale genomics.

Here, we develop *D*-plus, a maximum-likelihood (ML) framework that addresses these two confounding factors to improve the accuracy of introgression inference. Rather than assuming that a significant *D*-statistic reflects non-sister introgression, *D*-plus uses a hierarchical likelihood framework to evaluate alternative explanations for *ABBA–BABA* imbalance. Specifically, we first establish a species-tree model incorporating rate variation between sister lineages (M0; Fig. 1a) and then extends this framework to a suite of introgression models describing non-sister introgression, ghost introgression, and their co-occurrence (M1-M7; Fig. 1b–c). Through sequential likelihood-ratio tests (LRTs), *D*-plus first assesses whether introgression is required to explain significant *D*-statistics and subsequently identifies the type of supported introgression events (Fig. 2).

**Figure 1.**
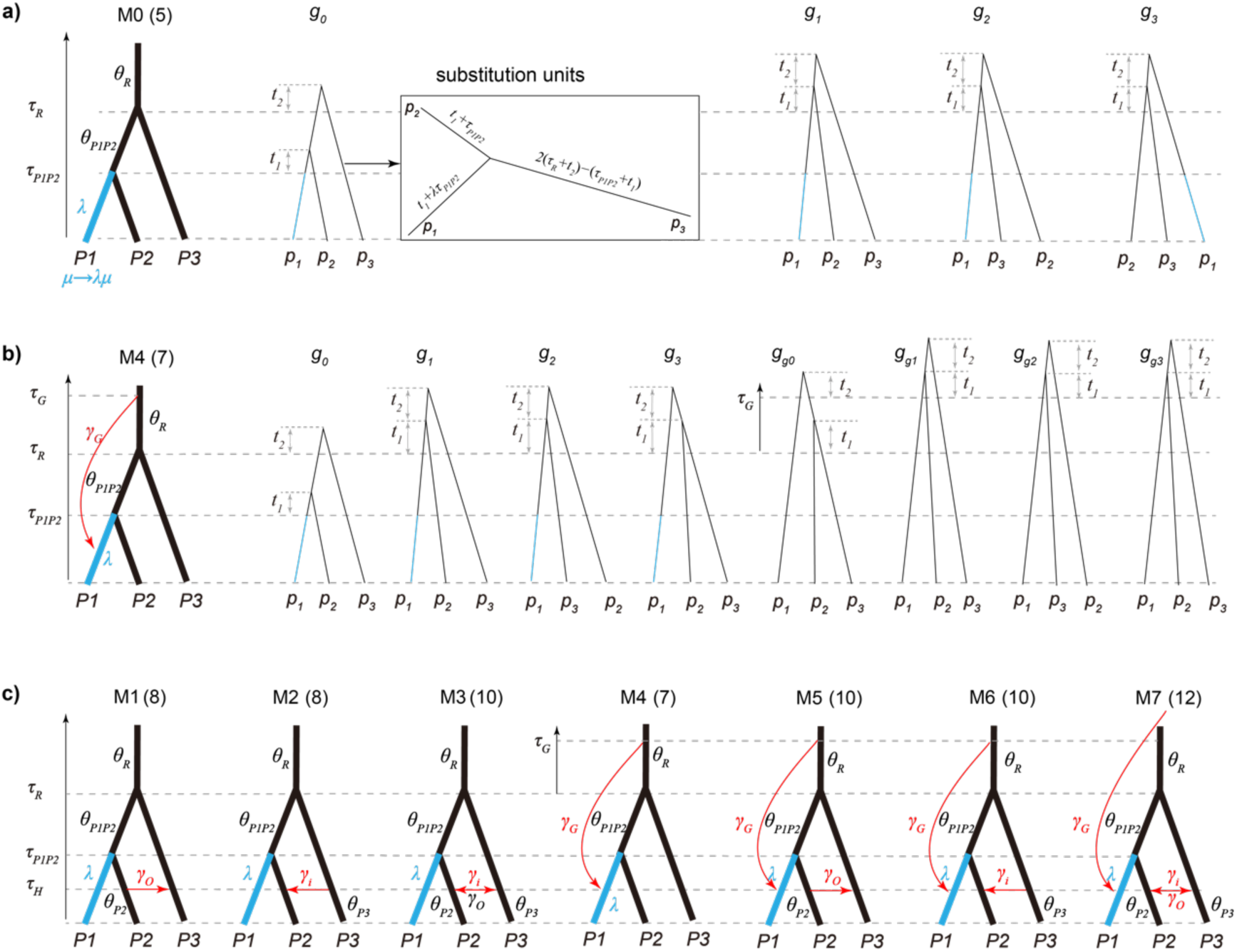
Models based on the species tree ((P1, P2), P3) incorporating rate variation without introgression (a) M0 or with introgression (b-c) M1-M7. Numbers within parentheses beside each model name indicate the total number of free parameters, including divergence and introgression times *τ*s, population-size parameters *θ*s, relative rate *λ* of the P1 lineage, and introgression probabilities *γ*s, as annotated on each schematic. For panels a-b, the right-hand side displays the set of feasible gene trees under the corresponding model shown on the left. Blue-highlighted segments along gene-tree branches experience substitution rate variation. Among introgression models, only M4 is illustrated with its full gene-tree repertoire here; detailed gene-tree configurations for all remaining introgression models are provided in Supplementary Text S1. Note that for all models involving ghost introgression, the ghost introgression event is assumed to occur at the present time.

**Figure 2.**
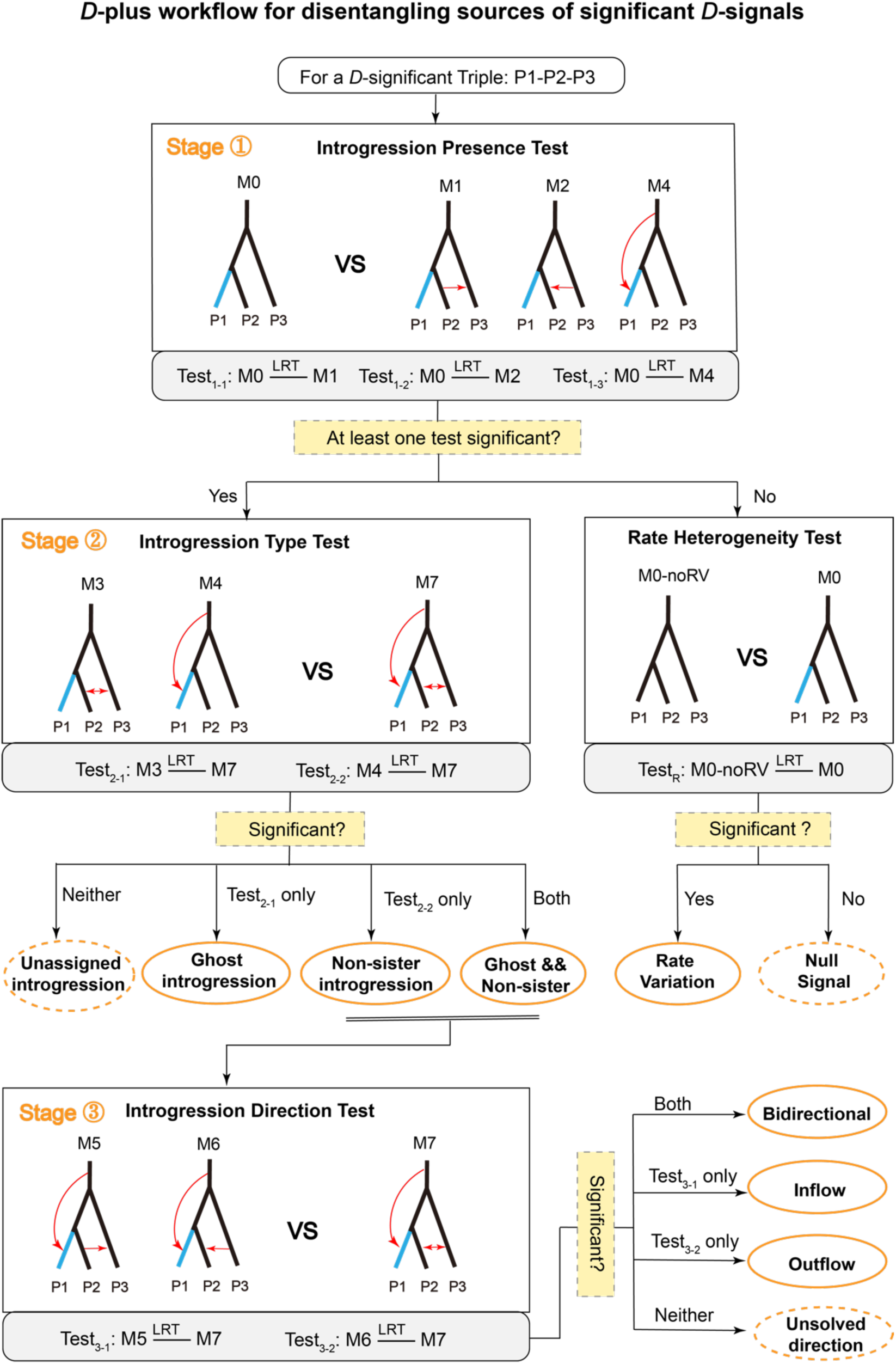
Workflow of *D*-plus.

The paper is structured as follows. First, we lay out the theoretical foundation of our model framework and describe the stepwise sequential likelihood ratio testing (LRT) procedure that forms the core of *D*-plus. Second, we report a comprehensive set of simulation benchmarks evaluating the performance of *D*-plus across both substitution rate heterogeneity and introgression scenarios, quantifying its specificity (false-positive control) and sensitivity (false-negative control). Finally, we demonstrate the practical application of *D*-plus using an empirical phylogenomic dataset from the plant genus *Jaltomata* (Solanaceae; Wu et al. 2018). Collectively, our simulation benchmarks and empirical validation establish D-plus as a robust, reliable, and computationally efficient tool for introgression inference in phylogenomic studies.

## Theory and Methods

### Null species-tree model accommodating rate variation (M0)

We focus on 3 ingroup species of a *D*-statistic quartet under a fixed species tree topology ((P1,P2),P3), with one sample per species {*p*_1_, *p*_2_, *p*_3_}. Building upon the classical multispecies coalescent (MSC; Rannala and Yang 2003) model, we propose Model 0 (M0; Fig. 1a), which additionally accommodates evolutionary rate variation between sister lineages. The model M0 is governed by 5 parameters: divergence times (*τ_R_*, *τ_P_*_1*P*2_), ancestral population sizes (*θ_R_*, *θ_P_*_1*P*2_), and the relative evolutionary rate *λ*. Here, *τ = Tμ* and *θ =* 4*Nμ*, where *T* is time in generations, *N* is the diploid population size, and *μ* is the ancestral substitution rate per site per generation. We assume that only lineage P1 experiences rate variation, with its substitution rate scaled to *λμ*.

The coalescent process follows the MSC framework (Fig. 1a). Tracing backward in time, if sequences *p*_1_ and *p*_2_ coalesced before the root time *τ_R_*, they form the gene tree *g*_0_. Otherwise, all 3 sequences enter the root population, where any pair would coalesce first with equal probability, producing one of the gene trees *g*_1_, *g*_2_, or *g*_3_. The joint probability density of these gene trees with coalescent times (*t*_1_, *t*_2_) is given by

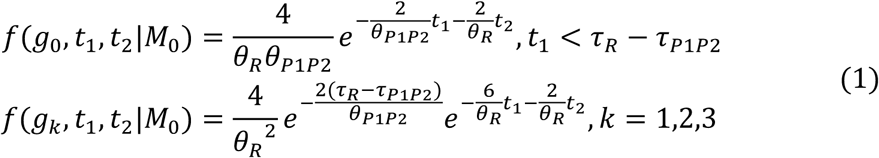

Crucially, the rate modification for lineage P1 carries over to all these gene trees: the portion of the *p*_1_ branch evolving within P1 lineage (highlighted in blue in Fig. 1a) receives a scaled branch-length of *λτ_P_*_1*P*2_.

Then we consider sequence evolution along gene trees under the Jukes–Cantor substitution model (Jukes and Cantor 1969). Leveraging the time-reversible property of this substitution model, we convert rooted gene-tree genealogies into their unrooted counterparts (Fig. 1a). The resulting unrooted-tree branch lengths—denoted *b_p_*_1_, *b_p_*_2_, and *b_p_*_3_ —can be directly obtained from species divergence times (*τ_R_*, *τ_P_*_1*P*2_) and the coalescent times (*t*_1_, *t*_2_). An exception is *b_p_*_1_, which inherits the *λ*-governed scaling effect described above, and its explicit form is given by:

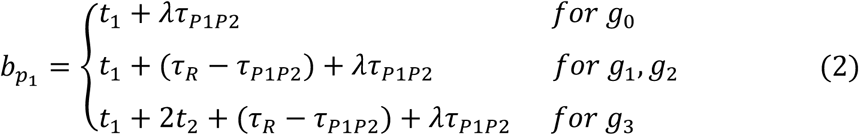

For observed sequence data, the alignment at each locus *i* can be summarized by a vector of site-pattern counts *D_i_* = {*n*_0_, *n*_1_, *n*_2_, *n*_3_, *n*_4_}, corresponding to the 5 distinct site-pattern categories: *xxx*, *yxx*, *xyx*, *xxy*, and *xyz*. Given an unrooted gene tree with specified branch lengths {*b_p_*_1_, *b_p_*_2_, *b_p_*_3_ }, the probabilities of the 5 site patterns at a single site are

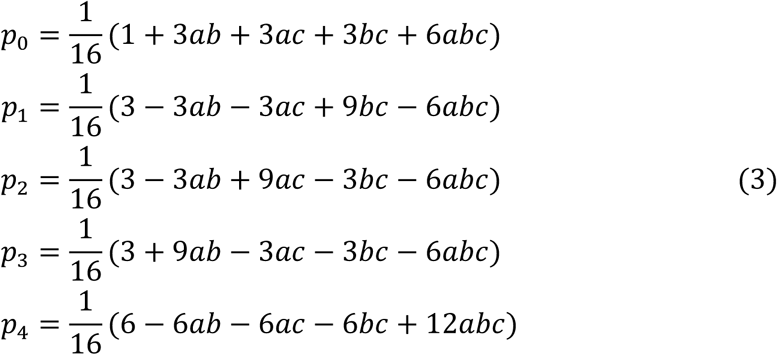

where 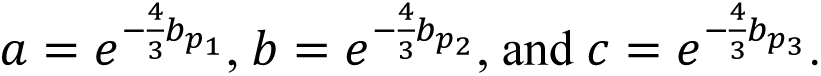 The probability of the observed count vector *D_i_* for locus *i* under this gene tree follows a multinomial distribution:

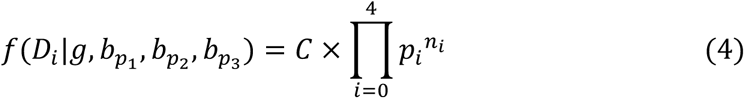

The probability of data *D_i_* under the model M_0_ is obtained by averaging over all possible gene trees and their associated coalescent times:

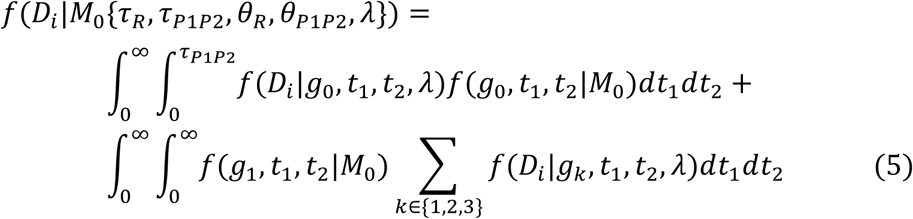

Finally, the log likelihood is sum over all the *L* loci

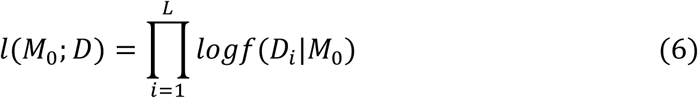

### Extended introgression models (M1-M7)

Starting from the baseline model M0, we further accommodate two types of introgression events: non-sister introgression and ghost introgression, both of which are typically associated with significant *D*-statistics (Tricou et al. 2022; Pang and Zhang 2024). Treating the two distinct directions of non-sister introgression — inflow (P3→P2) and outflow (P2→P3) — together with ghost introgression (ghost→P1) as three separable modelling units, we enumerate all valid combinations of these units — single, dual, or triple-event configurations — yielding 7 distinct introgression models denoted M1 to M7 (Fig. 1c). Here we take M4, which incorporates only a ghost introgression event, as an illustrative example (Fig. 1b); detailed descriptions of the remaining models are provided in Supplementary Text S1.

For the model M4, we introduce two additional parameters, *γ_G_* and *τ_G_*, to explicitly characterize the ghost introgression event, representing the introgression strength (i.e., the proportion of sequences introgressed) and the divergence time of the ghost lineage, respectively (Figure 1c). Two further assumptions are made: (1) the root population shares a single population size parameter, *θ_R_*, across both pre-and post-ghost divergence periods; and (2) the ghost introgression occurs at the present. The coalescent process follows the model of multispecies coalescent with introgression (MSci; Yu et al. 2014). Tracing backward in time, the coalescent process can be partitioned into 2 parts: (1) With probability 1 − *γ_G_*, sequence *p*_1_ enters the P1P2 ancestral branch and coalesces with sequence *p*_2_ prior to time *τ_R_*, yielding gene tree *g*_0_. Otherwise, all sequences enter the root, producing one of the gene trees *g*_1_, *g*_2_, or *g*_3_. (2) With probability *γ_G_*, sequence *p*_1_ enters the ghost branch. If sequences *p*_2_ and *p*_3_ coalesce before time *τ_G_*, this produces gene tree *g*_g0_. Otherwise, all sequences coalesce randomly after *τ_G_*, resulting in gene trees *g*_g1_, *g*_g2_, or *g*_g3_. The probability densities for the resulting gene trees are given by:

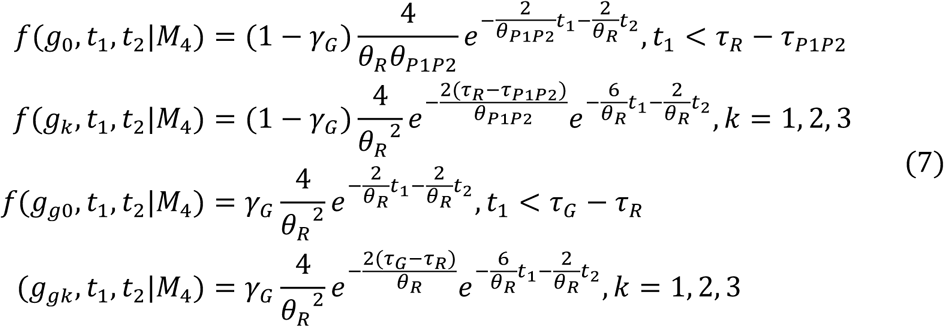

For unrooted counterparts of gene trees *g*_0_–*g*_3_, the branch length of branch *p*_1_ follows equation (2). For gene trees *g*_g0_*–g*_g3_, the branch length of branch *p*_1_ is not subject to rate variation in P1, and thus solely depends on species divergence times (*τ_R_*, *τ_G_*) and coalescent times (*t*_1_, *t*_2_). Finally, the probability of the observed data at locus *i* under model M4 is then:

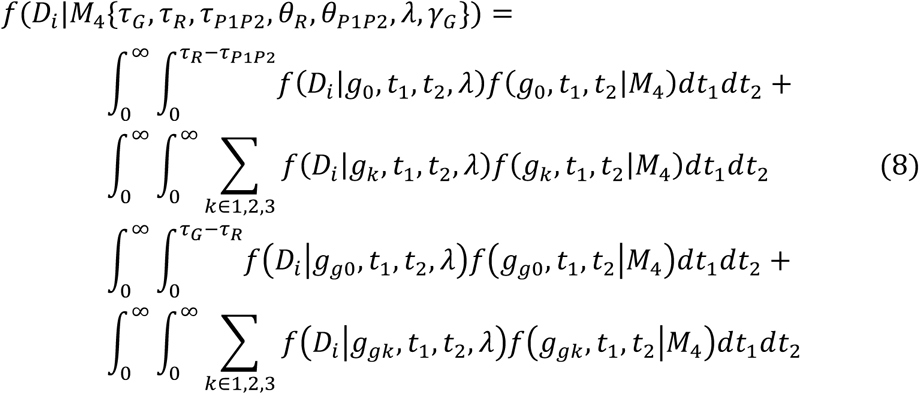

### Numerical Integration and ML Estimation

The *D*-plus program adopts the numerical procedures of the *3s* software (Yang 2010; Zhu and Yang 2012; Dalquen et al. 2017). For each model (M0-M7), the log-likelihood for data consisting of *L* loci entails at least 2*L* 2-D integrals over the two coalescent times (*t₁*, *t₂*) across gene tree classes. These integrals are computed via Gaussian quadrature, with a default of 16 quadrature points per dimension. Maximum likelihood estimates (MLE) were obtained using the BFGS quasi-Newton algorithm from the PAML package (Yang 2007). The program is implemented in C and supports multi-core parallel execution for per-locus log-likelihood computation.

### Likelihood Ratio Tests

For each statistically significant *D*-statistic triple, *D*-plus implements a three-stage sequential testing procedure (Fig. 2).

Stage 1: Introgression Presence Test. We first test for the genuine introgression signals, avoiding interference from rate variation. This step is achieved via three likelihood-ratio tests (LRT; Test_1-1_, Test_1-2_, and Test_1-3_; Fig. 2), in which the single-type introgression models M1, M2 (non-sister introgression with outflow or inflow direction), and M4 (ghost introgression) are each compared against the nested null model M0 (no introgression). Significant results from either LRT indicate the presence of introgression, triggering Stage 2 for type resolution. If neither LRT reaches significance, an ancillary rate-heterogeneity test is performed (see below).

Stage 2: Introgression Type Test. The full model M7, which accommodates both introgression types, is compared against each nested single-type model via LRT: Test_2-1_ contrasts M7 with the non-sister introgression model M3, thereby testing for the additional contribution of ghost introgression; Test_2-2_ contrasts M7 with the ghost introgression model M4, testing for the additional contribution of non-sister introgression. Together, these two contrasts establish whether the confirmed introgression signals arise from non-sister introgression, ghost introgression, or both.

Ancillary Test: Rate Heterogeneity Test. When Stage 1 yields no evidence of genuine introgression, we perform an LRT (Test_R_) for among-lineage rate heterogeneity, comparing M0 against the standard multispecies coalescent (MSC) model with no rate variation. This test assesses whether the significant *D*-statistic can be attributed to rate variation between sister lineages, a known confounder of *D*-statistic signals.

Stage 3: Introgression Direction Test. Conducted exclusively for cases with supported non-sister introgression from Stage 2, this step resolves its directionality via two LRTs (Test_3-1_ and Test_3-2_). The full model M7 is compared against each nested model that excludes one direction of non-sister introgression: M5 (ghost introgression + outflow) and M6 (ghost introgression + inflow). Each contrast tests whether the omitted direction is additionally required. Together, these two tests establish whether the recovered non-sister introgression proceeds as inflow, outflow, or bidirectional.

This sequential testing framework allowed us to rigorously identify the source of a significant *D* signal—whether it arises from ghost introgression, inflow, outflow (which may co-occur), or rate heterogeneity between sister lineages.

It is important to emphasize that the LRTs implemented in *D*-plus violate standard regularity conditions for the asymptotic *χ^2^* distribution. Under standard likelihood theory, the LRT statistic *2Δℓ* asymptotically follows a *χ^2^* distribution with degrees of freedom equal to the parameter-count difference between the two competing models. This large-sample approximation, however, depends on regularity assumptions that are violated for most of our tests. For instance, consider Test_1-3_ (M4 vs. M0): the null hypothesis M0 corresponds to fixing *γ_G_* at 0 in the alternative hypothesis M4, which lies on the boundary of the parameter space; additionally, when *γ_G_* = 0, the parameter *τ_G_* becomes unidentifiable in M4. As a result, the true null distribution deviates from the canonical *χ^2^*. Following Zhu and Yang (2012), we still adopt the *χ^2^* distribution for hypothesis testing at the 1 % significance level, as this approach has been shown to yield conservative test outcomes.

## Simulations

We performed extensive computer simulations to evaluate the type-I and type-II error properties of our implemented *D*-plus LRT framework and to quantify the precision of parameter estimation.

All simulations were conducted under a four-taxon species-tree framework (((*A*, *B*), *C*), *O*), with fixed divergence times of 10^6^ generations for lineage *C* and 3×10^6^ generations for the outgroup *O* (Fig. 3a and 4a). The effective population size (2*N*) was fixed at 10^5^ across all lineages, and the ancestral substitution rate per generation per site was set to *μ_R_* = 10⁻^8^. Within this basic framework, we further introduced different patterns of lineage-specific substitution-rate variation and introgression events to construct the specific simulation scenarios described below.

**Figure 3.**
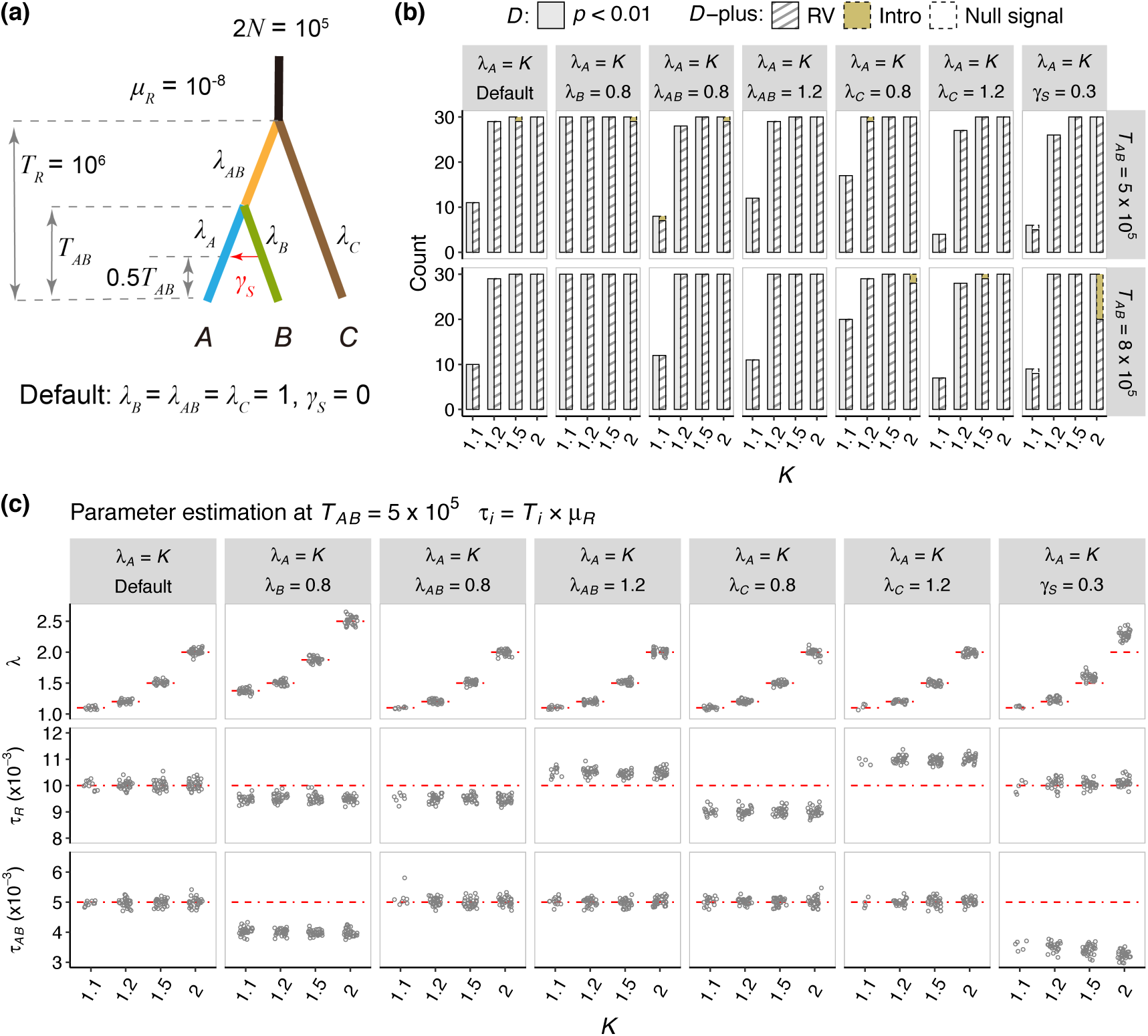
Performance of *D*-plus under various rate-variation scenarios. (a) Schematic and parameter settings of the four-taxon simulation (((*A*, *B*), *C*), *O*). The outgroup is omitted for simplicity, with its divergence time fixed at 3×10^6^ generations. *λᵢ* denotes the relative evolutionary rate of lineage *i* with respect to the root lineage *R*. Different rate-variation scenarios are labeled on the top strips in (b) and (c); key parameters *T_AB_* and *λ_A_* are shown on the right strips and x-axis, respectively. (b) Performance of *D*-plus in resolving the sources of significant *D* signals. For each condition on the x-axis, the left gray bar represents the number of replicates yielding significant *D* statistics (*p* < 0.01), while the right bar shows *D*-plus classification results, categorized into rate variation (RV), intro (introgression), or null signals (where neither rate variation nor introgression was detected). Notably, among the 4 replicates classified as introgression, 3 were identified as ghost introgression and 1 as unassigned introgression. (c) Parameter estimation at *T_AB_* = 5 × 10⁵, shown only for cases where *D*-plus successfully identified RV as the source. The red lines indicate the true parameter values; for *λ*, the red line represents the relative evolutionary rate of the sister lineages, computed as *λ_A_*/*λ_B_*. Gray points denote the estimated values. Results for *T_AB_* = 8 × 10⁵ are provided in Figure S2.

**Figure 4.**
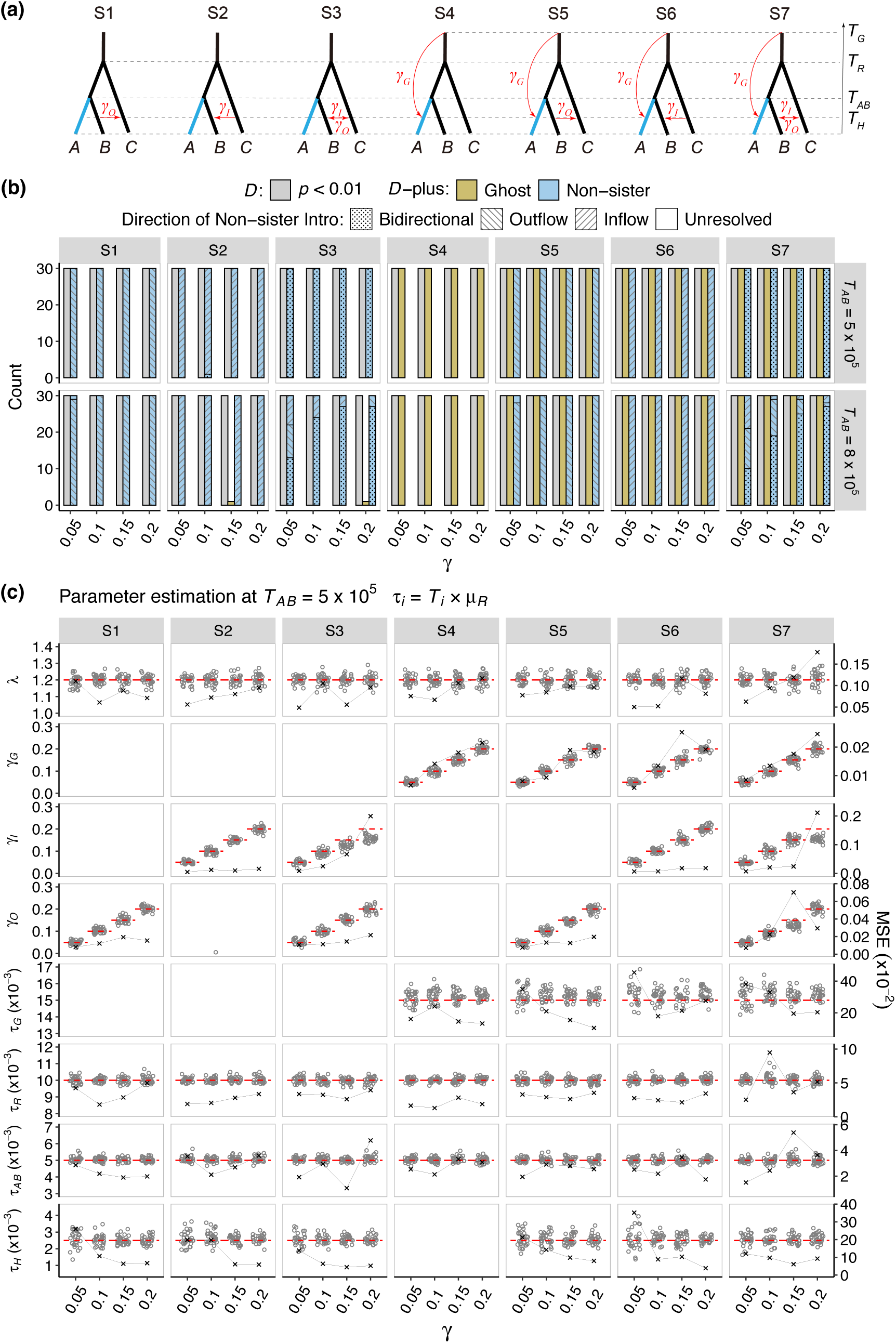
Performance of *D*-plus under various introgression scenarios. (a) Schematic illustration for 7 simulated introgression scenarios. *γ_G_*, *γ_I_*, and *γ*_O_ denote the introgression probabilities for ghost introgression, inflow and outflow events, respectively. The divergence times of the ghost lineage and lineage *C* are *T_G_* = 1.5×10^6^ and *T_R_* =10^6^ generations, with introgression time *T_H_* = 2.5×10^5^. Other parameters, including the outgroup, effective population size, and substitution rate, follow the settings in Figure 3. (b) Performance of *D*-plus in resolving the sources of significant *D* signals. For each condition on the x-axis, the leftmost gray bar represents the number of replicates with significant *D* statistics (*p* < 0.01), while the remaining bars show *D*-plus detection outcomes, which may correspond to ghost, non-sister, or unassigned introgression (i.e., significant introgression detected but not assigned to any specific type). Within non-sister introgression events, the fill patterns distinguish among directional categories: bidirectional, inflow, outflow, and unresolved (where neither direction could be confidently detected). (c) Parameter estimation from the inferred introgression model at *T_AB_* = 5 × 10⁵. Red lines indicate the true parameter values, and gray points denote the estimated values. The right-hand y-axis shows the mean squared error (MSE), defined as 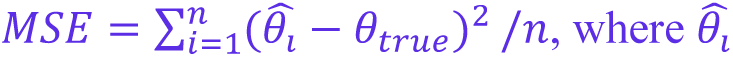 is the parameter estimate from replicate *i*, *θ_true_* is the true parameter value, and *n* is the total number of replicates. Results for *T_AB_* = 8 × 10⁵ are provided in Figure S3.

### Simulation 1: Type-I Error Evaluation

The first simulation suite accessed the type-I error (false-positive; FP) rate of *D*-plus under conditions in which rate variation was the sole cause of the *ABBA–BABA* imbalance. We established 7 distinct rate-variation scenarios (Fig. 3a), all involving rate variation in lineage *A* (*λ_A_*), combined with (1) null (default); (2) a slower substitution rate in the *B* clade (*λ_B_* = 0.8); (3) a slower rate in the *AB* ancestral clade (*λ_AB_* = 0.8); (4) a faster rate in the *AB* ancestral clade (*λ_AB_* = 1.2); (5) a slower rate in the *C* clade (*λ_C_* = 0.8); (6) a faster rate in the *C* clade (*λ_C_* = 1.2); and (7) sister introgression with introgression probability *γ_S_* = 0.3 and introgression time 0.5 × *T_AB_* (see below). For each scenario, we varied two key parameters: the divergence time between the sister lineages, *T_AB_* = 5 × 10⁵ or 8 × 10⁵ generations, and the relative rate of lineage *A*, *λ_A_* = 1.1, 1.2, 1.5, or 2.0.

Under each scenario and parameter combination, we simulated 30,000 independent gene trees with one sample per lineage using *fastsimcoal* v2.8 (Excoffier et al. 2013). Since *fastsimcoal2* does not natively implement lineage-specific substitution rate variation, we introduced rate heterogeneity via customized sampling time adjustments. We further generated 1,000-bp nucleotide sequences for each gene tree in *Seq-Gen* (Rambaut and Grass 1997) under the JC69 substitution model (Jukes and Cantor 1969). Each simulation condition was replicated 30 times. For downstream *D*-statistic analysis, per-gene sequences were concatenated to produce a total genome length of 30 Mb. Custom Python scripts were used to compute *D*-statistics and their standard errors (SE) via the block-jackknife resampling method. Statistical significance was determined using a z-test (z = D / SE(D)) with a *p*-value threshold of 0.01. Simulated data sets yielding significant *D*-signals were subsequently analyzed with *D*-plus using a subsample of 1,000 independent loci (excluding the outgroup *O*), unless otherwise specified. A single run of the *D*-plus implementation took 2–4 minutes per dataset.

The results are summarized in Figure 3 and Figure S2. Overall, the *D*-statistic was highly sensitive to rate-variation effects. A mere 10% substitution-rate difference between sister lineages triggers spurious introgression detections in 10%–60% of replicates, and doubling the rate variation to 20% pushes the FP rate to nearly 100% across every rate-variation scenario we tested. By contrast, *D*-plus performed exceptionally well: it maintained low rates of false introgression calls (i.e., Intro; Fig. 3b), while reliably attributing significant *D*-statistic signals to their true source, namely rate variation (i.e., RV). In all rate-variation scenarios, the FP rate remained near or below the nominal 1% significance level, confirming that the *χ²*-distribution test is conservative (Fig. 3b). Only one exception to this pattern was observed for the scenario incorporating sister introgression: *D*-plus misattributed significant *D*-signals to (ghost) introgression with a FP rate of 33% under *T_AB_* = 8 × 10⁵ and *λ_A_* = 2.

Regarding parameter estimation, *D*-plus provided accurate estimates of the relative rate (*λ*) of sister lineages across most scenarios (Fig. 3c and Fig. S2). One clear deviation arose in scenarios involving sister introgression: *λ* was systematically overestimated, and this upward bias intensified with larger *T_AB_* and *λ_A_* values. For the divergence-time parameters *τ_R_* and *τ_AB_*, inference remained accurate under the default scenario where rate variation was restricted exclusively to lineage *A*. By contrast, introducing rate variation in additional lineages violated model assumptions of *D*-plus and introduced bias into *τ* estimates. For example, in scenarios with *λ_B_* = 0.8, the estimated *τ_AB_* instead captures *T_AB_ × μ_B_*. Converting these scaled parameter estimates into absolute divergence times (in generations) using the ancestral mutation rate *μ_R_* therefore yields underestimations.

### Simulation 2: Type-II Error Evaluation

The second simulation suites quantified the type-II error (false-negative; FN) rate of *D*-plus under different introgression scenarios. Under the basic species-tree framework, we designed 7 introgression scenarios (S1-S7; Fig. 4a) covering all possible combinations of inflow, outflow, and ghost introgression events.

Introgression time were uniformly placed at *T_H_* = 2.5 × 10^5^ generations, and the ghost-lineage divergence time was fixed at *T_G_* = 1.5 × 10⁶ generations. Notably, *fastsimcoal2* does not support the bidirectional introgression model adopted by our *D*-plus method, which was originally proposed by Yang and Flouri (2022). We therefore specified an inflow event followed sequentially by an outflow event for scenarios S3 and S7. For each introgression scenario, we fixed *λ_A_* = 1.2, while varying the sister-lineage divergence time, *T_AB_* = 5 × 10⁵ or 8 × 10⁵ generations, and the introgression strength, *γ_I_*, *γ*_>_, *γ_G_* = 0.05, 0.1, 0.15, 0.2. Each *D*-plus analysis run required 15–30 minutes per dataset on a single CPU core.

The results are summarized in Figure 4 and Figure S3. The *D*-statistic flagged every single replicate as significant across all scenarios and parameter combinations (Fig. 4b). Remarkably, *D*-plus recapitulated this 100% detection power using only 1,000 loci, compared with 30,000 loci for the *D*-statistic (Fig. 4b). It further achieved accurate classification of introgression type, distinguishing between ghost introgression, non-sister introgression, and their co-occurrence across nearly all replicates. Only two replicates showed misclassifications in the full test set, arising in scenarios S2 and S3. In both instances, *D*-plus correctly identified non-sister introgression yet falsely inferred ghost introgression.

In contrast, the ability of *D*-plus to resolve the direction of non-sister introgression was comparatively sensitive to the sister-lineage divergence time *Tab* (Fig. 4b). At *Tab* = 5 × 10⁵ generations, introgression direction was correctly identified across all replicates, except for one replicate in scenario S2 that misclassified inflow as bidirectional introgression. However, when *Tab* increased to 8 × 10⁵ generations, while direction assignment accuracy remained close to 100% for unidirectional cases, performance declined markedly for the bidirectional introgression scenarios S3 and S7. For S3 and S7 under the low introgression strength of *γ* = 0.05, only approximately 30–40% of replicates correctly detected bidirectional gene flow; the remainder recovered only a single direction (inflow or outflow). As *γ* increased, direction assignment accuracy rose accordingly, reaching 90% at the moderate *γ* = 0.2.

We examined the accuracy and precision of parameter estimation. Overall, estimates of all parameters across replicates clustered around their true values, except for two parameter sets that exhibited systematic biases. The first bias, affecting *γ_I_* and *γ_O_*, was restricted exclusively to scenarios with bidirectional non-sister introgression (S3 and S7; Fig. 4c). This bias arises from a mismatch between our simulation design and the *D*-plus model for bidirectional introgression (see simulation details above). The second corresponds to systematic upward bias in *τ_G_* estimates, which originates from the *D*-plus model’s assumption that ghost introgression occurs at the present day — an unrealistic assumption that deviates from our simulation setup and thereby introduces directional error.

Furthermore, the precision (random error) of parameter estimation depended on *T_AB_* and *γ*. First, for parameters *γ*, *T_G_*, and *T_AB_*, the dispersion of estimates was substantially larger at *T_AB_* = 8 × 10⁵ generations than at *T_AB_* = 5 × 10⁵ (Figs. 4c and S3). The effects on these parameters are not straightforward. Notably, elevated ILS did not impact estimates of the root time *T_R_*, yet it reduced the precision of estimates for sister-lineage divergence time *T_AB_* and ghost-lineage divergence time *T_G_*. Second, for the time parameters *τ_H_* and *τ_G_*, estimation dispersion declined monotonically with rising introgression strength *γ*. This pattern is expected: information for these parameters is encoded within introgressed loci, so estimation precision deteriorates under weak introgression, where fewer informative loci are available.

### Analysis of *Jaltomata* Data

We applied *D*-plus to the published transcriptome dataset comprising 14 *Jaltomata* species and the outgroup *Solanum lycopersicum* (Wu et al. 2018). *Jaltomata* falls into major subgroups defined by mature fruit colour (Miller et al. 2011). Prior genomic analyses of this system uncovered abundant statistically significant *D*-statistic signals across species. Such signals were particularly prevalent when the early-diverging purple-fruited species were assigned as the P3 taxon, and these were interpreted as key evidence for recurrent non-sister introgression between the purple-fruited clade and other subgroups (Wu et al. 2018; Fig. 5a and Table 1). In addition, our earlier work (Pang et al. 2025), employing the relative rate test (hereinafter referred to as RR test; Graur and Li 2000), uncovered substantial lineage-specific substitution-rate variation among species pairs within *Jaltomata* (Fig. 5b-c). Given the co-occurrence of strong *D*-statistic signals and pronounced rate heterogeneity across the genus, we set out to: (1) disentangle whether significant *D*-statistics stem from rate-variation artefacts rather than genuine introgression; and (2) characterize well-supported introgressions, with a focus on determining their type.

**Figure 5.**
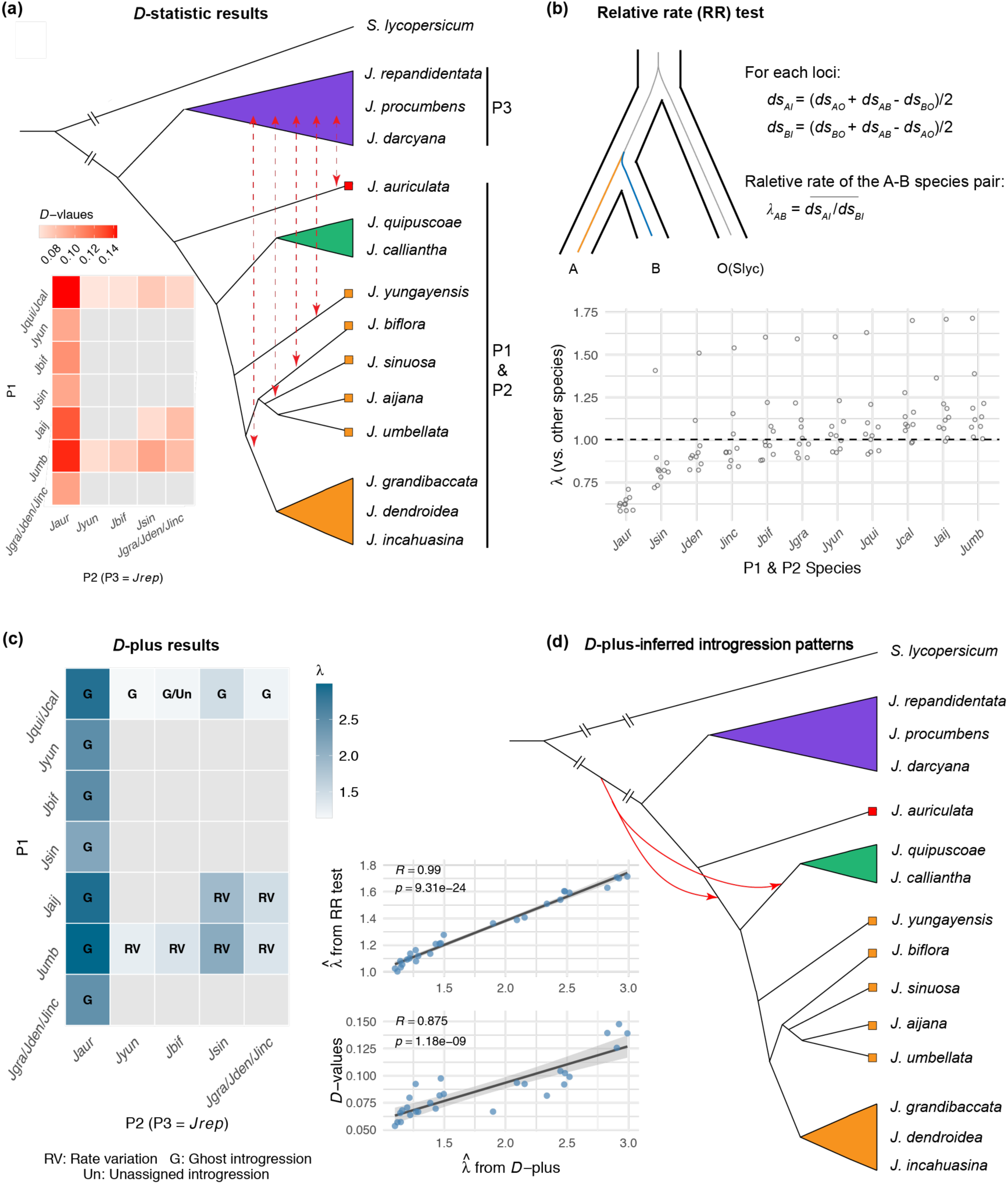
*D*-plus results for the *Jaltomata* group. (a) *D*-statistic results and previously inferred introgression patterns. The heatmap on the lower left provides detailed *D*-statistic results for all tested triples (P1–P2–P3), with P1 and P2 shown on the vertical and horizontal axes, respectively, and P3 fixed as *J. repandidentata* (*Jrep*) of the purple-fruited clade. For simplicity, species within the green-fruited clade (*Jqui/Jcal*) and orange-fruited clade (*Jgra/Jden/Jinc*) were each merged into a single analytical unit. Colored cells correspond to significant triples (*p* < 0.01), with color intensity encoding *D*-value magnitude—a single value per individual triple, and the mean across valid triples for merged analytical units at P1 or P2. The *Jaltomata* phylogeny at right (modified from Wu et al., 2018, Figure 3b) illustrates 5 inferred non-sister introgression events detected from *D*-statistic analyses, marked by red dashed lines. Tip colors of the phylogeny correspond to fruit color for each species. (b) Pronounced evolutionary-rate heterogeneity The upper schematic illustrates the methodological details of the relative-rate test. The lower scatter plot presents pairwise relative-rate estimates (*λ*) from this method for *Jaltomata*. Analyses are restricted to P1 and P2 lineages showing significant *D*-statistic signals. (c) Reanalyzed results from *D*-plus. The left heatmap presents detailed *D*-plus outputs for all *D*-significant trios. Each colored cell is annotated with the specific source of the *D*-signal identified by *D*-plus: RV, rate variation; G, ghost introgression; Un: unassigned introgression. The label “G/Un” denotes that the *Jqui*–*Jbif*–*Jrep* and *Jcal*–*Jbif*–*Jrep* trios produced distinct classifications, corresponding to “G” and “Un”, respectively. Color intensity scales with *λ* estimates for individual P1–P2 pairs, and with mean *λ* values across multiple pairs for cell involving merged analytical units. The lower right panel shows correlation plots: top, comparison of *λ* estimates derived from *D*-plus and the relative-rate test; bottom, correlation between *λ* estimates inferred by *D*-plus and *D*-values. Pearson’s correlation coefficients (*R*) measure the strength and direction of linear association between the paired metrics, with associated *p*-values testing the null hypothesis of no linear correlation. (d) Two introgression events inferred from the *D*-plus outputs.

**Table 1.** Significant D-statistics for Jaltomata triples and corresponding D-plus analyses: identification of D-signal sources and parameter estimation.

| Triple | *D*-statistic | *D*-plus | $\tau_R$ | $\tau_{P1P2}$ | $\tau_G$ | $\gamma_G$ | $\lambda$ |
| --- | --- | --- | --- | --- | --- | --- | --- |
| *Jqui*–*Jaur*–*Jrep* | 0.139486 | G | 0.00274 | 0.00088 | 0.00515 | 0.0636 | 2.8280 |
| *Jcal*–*Jaur*–*Jrep* | 0.147689 | G | 0.00274 | 0.00087 | 0.00528 | 0.0604 | 2.9229 |
| *Jyun*–*Jaur*–*Jrep* | 0.091996 | G | 0.00274 | 0.00091 | 0.00517 | 0.0673 | 2.4767 |
| *Jbif*–*Jaur*–*Jrep* | 0.102476 | G | 0.00272 | 0.00091 | 0.00486 | 0.0686 | 2.4834 |
| *Jsin*–*Jaur*–*Jrep* | 0.092491 | G | 0.00274 | 0.00087 | 0.00540 | 0.0534 | 2.1547 |
| *Jaij*–*Jaur*–*Jrep* | 0.125825 | G | 0.00274 | 0.00091 | 0.00510 | 0.0660 | 2.9050 |
| *Jumb*–*Jaur*–*Jrep* | 0.139189 | G | 0.00275 | 0.00090 | 0.00527 | 0.0643 | 2.9907 |
| *Jgra*–*Jaur*–*Jrep* | 0.099047 | G | 0.00273 | 0.00092 | 0.00515 | 0.0694 | 2.5201 |
| *Jden*–*Jaur*–*Jrep* | 0.081649 | G | 0.00272 | 0.00092 | 0.00505 | 0.0606 | 2.3356 |
| *Jinc*–*Jaur*–*Jrep* | 0.104418 | G | 0.00273 | 0.00089 | 0.00498 | 0.0675 | 2.4462 |
| *Jqui*–*Jyun*–*Jrep* | 0.057092 | G | 0.00353 | 0.00198 | 0.00573 | 0.0300 | 1.1200 |
| *Jcal*–*Jyun*–*Jrep* | 0.068167 | G | 0.00352 | 0.00196 | 0.00577 | 0.0287 | 1.1593 |
| *Jumb*–*Jyun*–*Jrep* | 0.066864 | RV | 0.00344 | 0.00159 | NA | NA | 1.2715 |
| *Jqui*–*Jbif*–*Jrep* | 0.057318 | G | 0.00353 | 0.00179 | 0.00668 | 0.0139 | 1.1474 |
| *Jcal*–*Jbif*–*Jrep* | 0.070717 | Un | 0.00352 | 0.00177 | NA | NA | 1.1994 |
| *Jumb*–*Jbif*–*Jrep* | 0.074898 | RV | 0.00339 | 0.00121 | NA | NA | 1.3874 |
| *Jqui*–*Jsin*–*Jrep* | 0.069697 | G | 0.00333 | 0.00138 | 0.00677 | 0.0112 | 1.4310 |
| *Jcal*–*Jsin*–*Jrep* | 0.083144 | G | 0.00332 | 0.00136 | 0.00701 | 0.0107 | 1.4973 |
| *Jumb*–*Jsin*–*Jrep* | 0.093479 | RV | 0.00323 | 0.00072 | NA | NA | 2.0918 |
| *Jaij-Jsin-Jrep* | 0.066922 | RV | 0.00321 | 0.00079 | NA | NA | 1.8981 |
| *Jqui-Jden-Jrep* | 0.079758 | G | 0.00345 | 0.00172 | 0.00671 | 0.0145 | 1.2175 |
| *Jcal-Jden-Jrep* | 0.092449 | G | 0.00343 | 0.00170 | 0.00650 | 0.0162 | 1.2677 |
| *Jumb-Jden-Jrep* | 0.097561 | RV | 0.00334 | 0.00121 | NA | NA | 1.4733 |
| *Jaij-Jden-Jrep* | 0.081855 | RV | 0.00331 | 0.00109 | NA | NA | 1.4629 |
| *Jqui-Jgra-Jrep* | 0.053666 | G | 0.00358 | 0.00189 | 0.00643 | 0.0166 | 1.1001 |
| *Jcal-Jgra-Jrep* | 0.065687 | G | 0.00359 | 0.00183 | 0.00645 | 0.0141 | 1.1432 |
| *Jumb-Jgra-Jrep* | 0.066778 | RV | 0.00346 | 0.00136 | NA | NA | 1.2889 |
| *Jcal-Jinc-Jrep* | 0.063820 | G | 0.00348 | 0.00172 | 0.00646 | 0.0151 | 1.2237 |

We re-evaluated all triples that yielded significant *D*-statistics, where the purple-fruited species *J. repandidentata* was set as the P3 taxon. This re-analysis was performed with *D*-plus, using a total of 6,431 one-to-one orthologous alignments. For each triple, *D*-plus was run with 3 independent repeats (*repeat* = 3) to avoid convergence to local optima, completing in ∼30 minutes with 10 threads. All results are summarized in Figure 5 and Table 1.

First, for 7 *D*-significant triples featuring *J. umbellata* or *J. aijana* as P1 and other orange-fruited lineages as P2, *D*-plus attributed these *D*-signals entirely to rate-variation artifacts rather than biological introgression (Fig. 5c, left panel); *D*-plus inferred that *J. umbellata* and *J. aijana* exhibited evolutionary rates 27 %–109 % higher than those of other orange-fruited lineages (*λ* = 1.27–2.09; Table 1).

Second, across 10 *D*-significant triples with red-fruited *J.auriculata* as P1 and green-or orange-fruited lineages as P2, *D*-plus consistently detected introgression signals, all of which were classified as ghost introgression (Fig. 5c). Estimates of ghost-donor divergence time (*τ_G_* = 0.00486–0.00517; Table 1) and introgression probability (*γ_G_* = 0.053–0.067; Table 1) were tightly clustered across these triples. This consistent pattern supports a single ghost introgression event from an ancestral *Jaltomata* lineage into the common ancestor of the green-and orange-fruited clades (Fig. 5d). This finding aligns with SNaQ inferences (Tiley et al., 2023), which are robust to rate variation through exclusive reliance on gene-tree topologies, yet conflicts with our earlier BPP-based analyses (Pang and Zhang 2024) favouring an outflow event. This discrepancy likely arises because the full-likelihood BPP framework does not account for rate variation and is thus prone to associated biases. Indeed, substantial rate heterogeneity was inferred between *J. auriculata* and its respective P2 lineages (*λ* = 2.155–2.991; Table 1).

Third, the remaining 11 significant triples with the green-fruited lineages as P1 and orange-fruited lineages as P2 also exhibited ghost introgression signals in *D*-plus, with the exception of one trio (*J. calliantha*–*J. biflora*–*J. repandidentata*) in which introgression was confirmed but its type remained unassigned. Under parsimonious reasoning, these patterns imply a shared ghost introgression shaping the green-fruited clade (Fig. 5d).

Finally, we compared relative-rate estimates *λ* from *D*-plus with those from the RR test. Interestingly, the two sets of estimates exhibited strong linear correlation (R = 0.99 and *p* = 3.11e-14; Fig. 5c), but *D*-plus yielded systematically larger *λ* estimates (*λ*^T^ *=* 1.100–2.991; Table 1) than the RR test (*λ*^T^ *=* 1.003–1.714). This discrepancy likely arises from two factors. First, the RR test assumes that lineages coalesce immediately upon entering the ancestral population (i.e., no deep coalescence), an oversimplification that inherently underestimates the magnitude of rate variation. Second, potential sister introgression may inflate *λ* estimates from *D*-plus. Besides, *D*-plus-inferred *λ* values also showed a strong positive correlation with *D*-statistic values (*R* = 0.86 and *p* = 9.64e-6). This indicates that much of the variation in *ABBA-BABA* imbalance magnitude across trios in the *Jaltomata* system can be attributed to lineage-specific rate variation, rather than purely reflecting the extent of introgression.

## Discussion

### Robust false-positive control under substitution-rate heterogeneity

A persistent challenge in *D*-statistic-based introgression screening is that lineage-specific shifts in substitution rate can generate *ABBA–BABA* imbalance mimicking the signature of gene flow and yields pervasive false positives (Frankel and Ané 2023; Pang et al. 2025). Our simulations corroborate this well-recognized vulnerability: the *D*-statistic is highly sensitive to rate heterogeneity, producing significant signals with as little as 10 % rate variation between sister lineages even in the absence of introgression (Fig. 3).

In stark contrast, *D*-plus effectively decouples rate variation from introgression. It maintains low false-positive rates for introgression calls and correctly attributes significant *D*-signals to rate-heterogeneity artefacts (Fig. 3). This robust performance generalizes across a broad parameter space, holding for diverse lineage configurations involving rate variation, variable magnitudes of rate heterogeneity, and differing depths of sister-lineage divergence.

That said, there exists one pathological parameter regime in which *D*-plus misattributes *D*-signals to introgression. When sister introgression—an event that by itself does not produce an *ABBA–BABA* skew—co-occurs with substitution-rate heterogeneity, *D*-plus misinterprets the rate-variation-driven *D*-signal as evidence of ghost introgression. This failure mode is restricted to scenarios of rapid radiative branching (*T_AB_* = 8 × 10⁵ relative to *T_R_* = 10^6^) coupled with pronounced rate disparity (*λ* = 2) (Fig. 3b). Reassuringly, such extreme parameter combinations would be expected to be rare in natural systems, and thus are unlikely to distort inference in most empirical studies.

### Power and limitations for introgression detection and characterization

Beyond its strong false-positive control, *D*-plus achieves high sensitivity for both introgression detection and type identification. Across the full range of simulated introgression scenarios, *D*-plus retains 100 % statistical power to call introgression, while also accurately resolving introgression events into ghost introgression, non-sister introgression, or their simultaneous co-occurrence (Fig. 4b). To our knowledge, this ability to distinguish ghost introgression and non-sister introgression has previously only been achievable with the Bayesian method BPP (Cheng et al. 2026; Yang et al. 2026). By contrast, *D*-plus, as a maximum-likelihood method, delivers higher computational efficiency.

*D*-plus also delivers characterization of additional key features of introgression events, including direction, strength, and timing. We find that inference of both direction and strength is strongly influenced by internal branch lengths in the species tree, with greater reliability at longer internal branches (Fig. 4c). Under rapid radiative branching (*Tab* = 8 × 10⁵ relative to *Tr* = 10^6^), estimates of introgression strength exhibit greater stochastic error across all introgression scenarios. Direction inference is particularly challenging under weak bidirectional introgression, for which *D*-plus always detects only one of the two directions (i.e., inflow or outflow) (Fig. 4b). This limitation is theoretically expected: under a star-like topology, introgression direction is inherently non-identifiable when only a single sequence is sampled per species (Thawornwattana et al. 2023). We therefore recommend using more loci, particularly when applying *D*-plus to rapidly radiating systems, to provide sufficient information for reliable inference.

### Quantification of lineage-specific substitution-rate heterogeneity

Beyond its core role in introgression inference, *D*-plus also quantifies lineage-specific substitution-rate variation—an evolutionary feature we have thus far treated solely as a confounding factor, but which is itself a fundamental, widely studied phenomenon in molecular evolution. As part of its standard analytical workflow, *D*-plus estimates the relative substitution rate (*λ*) between sister lineages. Our simulations show that these *λ* estimates are generally accurate across the parameter regimes examined, with the notable exception of systematic overestimation in the presence of sister introgression (Fig. 3c and 4c). Accurate quantification of rate heterogeneity can support a broad range of evolutionary research applications. For example, lineage-specific substitution rates can be calibrated against empirically established rates from closely related model organisms, enabling rate estimation in non-model clades for which experimental estimates are unavailable.

### Empirical insights from the Jaltomata system

Our empirical reanalysis of the *Jaltomata* system demonstrates the practical utility of *D*-plus in disentangling the distinct sources of *D*-statistic signals in a real-world biological system. First, *D*-plus effectively corrects bias arising from substitution-rate heterogeneity, revealing that several significant *D*-signals conventionally interpreted as evidence of introgression can instead be explained entirely by lineage-specific rate variation. Second, among the remaining *D*-significant triples, *D*-plus identifies two distinct ghost introgression events: one involving the common ancestor of the green-and orange-fruited lineages and another restricted to the green-fruited clade (Fig. 5d). This result challenges the previous interpretation of these signals as evidence of non-sister introgression. Third, *D*-plus reveals substantial substitution-rate heterogeneity across *Jaltomata* lineages, with relative rates (*λ*) ranging from 1.1-to 3-fold. Notably, *λ* estimates across trios are strongly positively correlated with *D*-statistic values, indicating that much of the variation in the magnitude of *ABBA–BABA* imbalance among trios in *Jaltomata* can be attributed to substitution-rate variation among species rather than the extent of introgression alone.

The *Jaltomata* case highlights that both substitution-rate heterogeneity and ghost introgression may be more widespread than commonly recognized. Accordingly, treating significant *D*-statistics as evidence of non-sister introgression by default carries a substantial risk of erroneous biological interpretation. We therefore recommend *D*-plus as a follow-up analysis whenever significant *D*-statistics are detected, allowing the sources of *ABBA–BABA* imbalance to be disentangled and more reliable introgression inferences to be obtained.

### Limitations and Future directions

Like all model-based coalescent frameworks, *D*-plus is built on a set of simplifying assumptions that define its current analytical scope. First, the current implementation uses a single sample per species. Future iterations of the framework could be designed to natively support multiple samples per lineage; incorporating these additional samples would leverage within-species genetic variation to substantially boost statistical power. This enhancement would be particularly impactful for resolving the direction of non-sister introgression in challenging radiative divergence regimes (Thawornwattana et al. 2023), where the current implementation performs poorly. Second, *D*-plus does not accommodate introgression between sister lineages. This process can distort branch-length distributions, causing misattribution to ghost introgression under certain conditions and introducing systematic bias into relative-rate estimation (Fig. 3). Extending the null model to incorporate sister introgression would mitigate these biased inferences, albeit with greater model complexity and higher computational overhead. Third, as with all full-likelihood coalescent methods, *D*-plus assumes no intra-locus recombination and free inter-locus recombination. Simulation evidence (Zhu et al. 2022; Thawornwattana et al. 2026) shows introgression testing remains robust to realistic recombination levels, with minimal bias in introgression probability estimates, making this a well-justified simplification.

## Supplementary Material

Supplementary data are available at….

## Author Contributions

X.-X.P. conceived and designed the study, implemented the software, and performed all simulations and empirical analyses. D.-Y. Z. provided key suggestions for simulations.

## Data Availability Statement

Simulated and empirical datasets are available in Zenodo at….Source code is publicly available at https://github.com/pangxiaoxu-123/D-plus.

## Funding Data Availability Statement

This work was supported by the National Natural Science Foundation of China (grant numbers 32600549); The Big Data Computing Platform for Western Ecological Environment and Regional Development and the Supercomputing Center of Lanzhou University.

