## supplemental materials for "*D*-plus: Beyond the *D*-statistic — Accurate Detection and Characterization of Introgression Despite Among-Lineage Rate Variation"

Supplementary Text S1

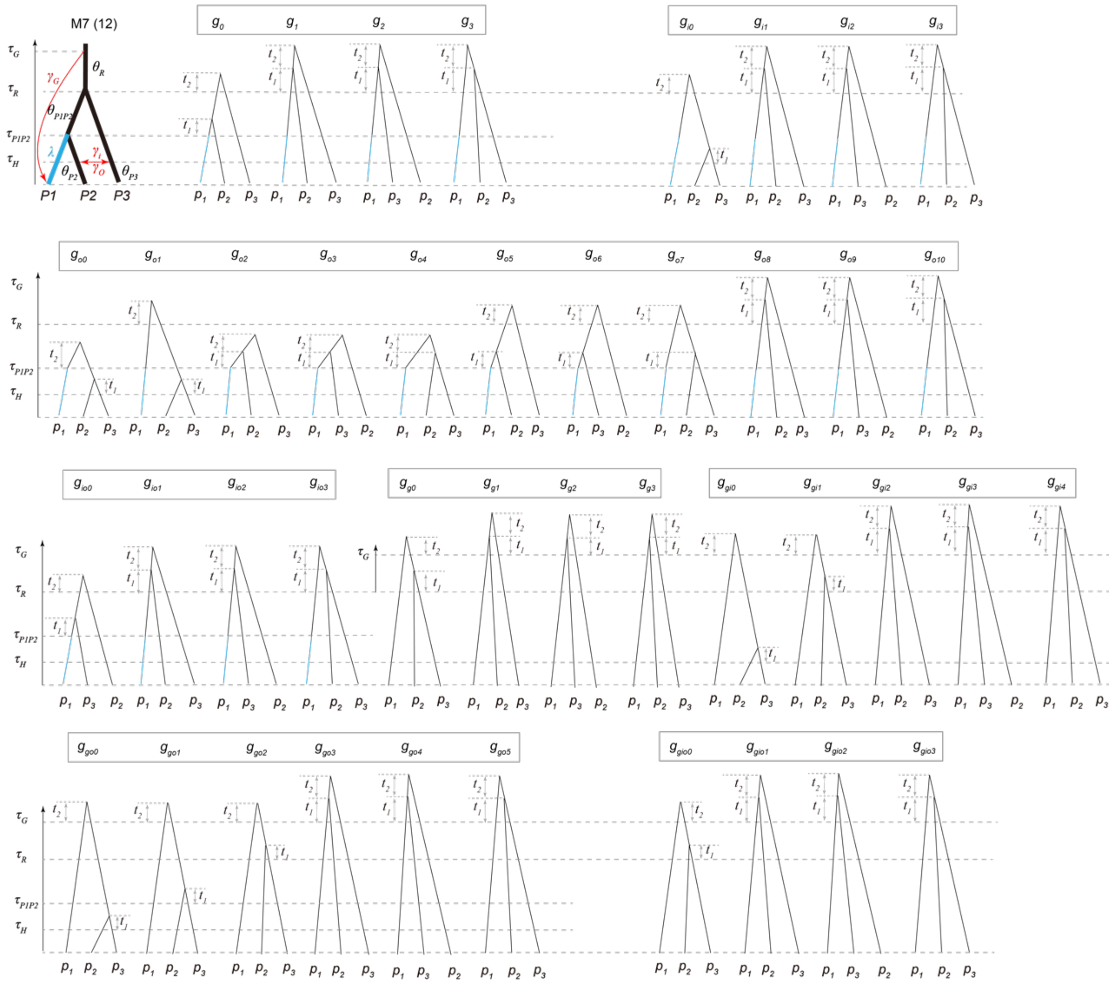

Figure S1. The set of feasible gene-tree topologies expected under the model M7.

Here, we derive the gene-tree probability distributions for our introgression models, starting with the most complex general model M7. Model M7 incorporates three distinct introgression events: ghost introgression, inflow, and outflow, governed by introgression probabilities $\gamma_G$, $\gamma_I$, and $\gamma_O$, respectively. When tracing the coalescent process backward in time, 8 mutually exclusive scenarios emerge, corresponding to all combinations of independent binary choices for each of the three sequences: each sequence may either follow an introgression path (red-labelled branches) or the standard species-tree path. These scenarios are defined as follows (Figure S1):

1. All 3 sequences follow the species-tree path, with probability $(1-\gamma_G)(1-\gamma_I)(1-\gamma_O)$, producing gene trees $g_0$–$g_3$;
2. Only sequence $p_2$ takes the introgression path, while $p_1$ and $p_3$ follow the standard path, with probability $(1-\gamma_G)\gamma_I(1-\gamma_O)$, producing gene trees $g_{i0}$–$g_{i3}$;

3. Only sequence $p_3$ takes the introgression path, while $p_1$ and $p_2$ follow the tree path, with probability $(1-\gamma_G)(1-\gamma_I)\gamma_O$, producing gene trees $g_{o0}$–$g_{o3}$;

4. Sequences $p_2$ and $p_3$ take their respective introgression paths, and $p_1$ follows the tree path, with probability $(1-\gamma_G)\gamma_I\gamma_O$, producing gene trees $g_{io0}$–$g_{io3}$;

5. Only sequence $p_1$ takes the ghost-introgression path, while $p_2$ and $p_3$ follow the tree path, with probability $\gamma_G(1-\gamma_I)(1-\gamma_O)$, producing gene trees $g_{g0}$–$g_{g3}$;

6. Sequences $p_1$ and $p_2$ take introgression paths, and $p_3$ follows the tree path, with probability $\gamma_G\gamma_I(1-\gamma_O)$, producing gene trees $g_{gi0}$–$g_{gi3}$;

7. Sequences $p_1$ and $p_3$ take introgression paths, and $p_2$ follows the tree path, with probability $\gamma_G(1-\gamma_I)\gamma_O$, producing gene trees $g_{go0}$–$g_{go3}$;

8. All three sequences take their respective introgression paths, with probability $\gamma_G\gamma_I\gamma_O$, producing gene trees $g_{gio0}$–$g_{gio3}$;

The probability densities for each of these resulting gene-tree classes are given by equation S1.

For the simpler introgression models M1-M6, the same gene-tree probability density equation (Equation S1) applies; their corresponding gene-tree probability expressions are obtained simply by fixing the introgression-probability parameters corresponding to omitted introgression events to zero.

$$
\begin{aligned}
f(g_0,t_1,t_2|M_7) &= (1-\gamma_G)(1-\gamma_I)(1-\gamma_O)\frac{4}{\theta_R\theta_{P1P2}}e^{-\frac{2}{\theta_{P1P2}}t_1-\frac{2}{\theta_R}t_2}, t_1<\tau_R-\tau_{P1P2}\\
f(g_k,t_1,t_2|M_7) &= (1-\gamma_G)(1-\gamma_I)(1-\gamma_O)\frac{4}{{\theta_R}^2}e^{-\frac{2(\tau_R-\tau_{P1P2})}{\theta_{P1P2}}}e^{-\frac{6}{\theta_R}t_1-\frac{2}{\theta_R}t_2}, k=1,2,3\\
f(g_{i0},t_1,t_2|M_7) &= (1-\gamma_G)\gamma_I(1-\gamma_O)\frac{4}{\theta_R\theta_{P3}}e^{-\frac{2}{\theta_{P3}}t_1-\frac{2}{\theta_R}t_2}, t_1<\tau_R-\tau_H\\
f(g_{ik},t_1,t_2|M_7) &= (1-\gamma_G)\gamma_I(1-\gamma_O)\frac{4}{{\theta_R}^2}e^{-\frac{2(\tau_R-\tau_H)}{\theta_{P3}}}e^{-\frac{6}{\theta_R}t_1-\frac{2}{\theta_R}t_2}, k=1,2,3\\
f(g_{o0},t_1,t_2|M_7) &= (1-\gamma_G)(1-\gamma_I)\gamma_O\frac{4}{\theta_{P1P2}\theta_{P2}}e^{-\frac{2}{\theta_{P2}}t_1-\frac{2}{\theta_{P1P2}}t_2}, t_1<\tau_{P1P2}-\tau_H, t_2<\tau_R-\tau_{P1P2}\\
f(g_{o1},t_1,t_2|M_7) &= (1-\gamma_G)(1-\gamma_I)\gamma_O\frac{4}{\theta_R\theta_{P2}}e^{-\frac{2(\tau_R-\tau_{P1P2})}{\theta_{P1P2}}}e^{-\frac{2}{\theta_{P2}}t_1-\frac{2}{\theta_R}t_2}, t_1<\tau_{P1P2}-\tau_H\\
f(g_{ok},t_1,t_2|M_7) &= (1-\gamma_G)(1-\gamma_I)\gamma_O\frac{4}{{\theta_{P1P2}}^2}e^{-\frac{2(\tau_{P1P2}-\tau_H)}{\theta_{P2}}}e^{-\frac{6}{\theta_{P1P2}}t_1-\frac{2}{\theta_{P1P2}}t_2}, t_1<\tau_R-\tau_{P1P2}, t_2<\tau_R-\tau_{P1P2}-t_1, k=2,3,4\\
f(g_{ok},t_1,t_2|M_7) &= (1-\gamma_G)(1-\gamma_I)\gamma_O\frac{4}{\theta_R\theta_{P1P2}}e^{-\frac{2(\tau_{P1P2}-\tau_H)}{\theta_{P2}}-\frac{2(\tau_R-\tau_{P1P2}-t_1)}{\theta_{P1P2}}}e^{-\frac{6}{\theta_{P1P2}}t_1-\frac{2}{\theta_R}t_2}, t_1<\tau_R-\tau_{P1P2}, k=5,6,7\\
f(g_{ok},t_1,t_2|M_7) &= (1-\gamma_G)(1-\gamma_I)\gamma_O\frac{4}{{\theta_R}^2}e^{-\frac{2(\tau_{P1P2}-\tau_H)}{\theta_{P2}}-\frac{6(\tau_R-\tau_{P1P2})}{\theta_{P1P2}}+t_1}e^{-\frac{6}{\theta_R}t_1-\frac{2}{\theta_R}t_2}, k=8,9,10\\
f(g_{io0},t_1,t_2|M_7) &= (1-\gamma_G)\gamma_I\gamma_O\frac{4}{\theta_R\theta_{P1P2}}e^{-\frac{2}{\theta_{P1P2}}t_1-\frac{2}{\theta_R}t_2}, t_1<\tau_R-\tau_{P1P2}\\
f(g_{iok},t_1,t_2|M_7) &= (1-\gamma_G)\gamma_I\gamma_O\frac{4}{{\theta_R}^2}e^{-\frac{2(\tau_R-\tau_{P1P2})}{\theta_{P1P2}}}e^{-\frac{6}{\theta_R}t_1-\frac{2}{\theta_R}t_2}, k=1,2,3\\
f(g_{g0},t_1,t_2|M_7) &= \gamma_G(1-\gamma_I)(1-\gamma_O)\frac{4}{{\theta_R}^2}e^{-\frac{2}{\theta_R}t_1-\frac{2}{\theta_R}t_2}, t_1<\tau_G-\tau_R\\
f(g_{gk},t_1,t_2|M_7) &= \gamma_G(1-\gamma_I)(1-\gamma_O)\frac{4}{{\theta_R}^2}e^{-\frac{2(\tau_G-\tau_R)}{\theta_R}}e^{-\frac{6}{\theta_R}t_1-\frac{2}{\theta_R}t_2}, k=1,2,3\\
f(g_{gi0},t_1,t_2|M_7) &= \gamma_G\gamma_I(1-\gamma_O)\frac{4}{\theta_R\theta_{P3}}e^{-\frac{2}{\theta_{P3}}t_1-\frac{2}{\theta_R}t_2}, t_1<\tau_R-\tau_H\\
f(g_{gi1},t_1,t_2|M_7) &= \gamma_G\gamma_I(1-\gamma_O)\frac{4}{{\theta_R}^2}e^{-\frac{2(\tau_R-\tau_H)}{\theta_{P3}}}e^{-\frac{2}{\theta_R}t_1-\frac{2}{\theta_R}t_2}, t_1<\tau_G-\tau_R\\
f(g_{gik},t_1,t_2|M_7) &= \gamma_G\gamma_I(1-\gamma_O)\frac{4}{{\theta_R}^2}e^{-\frac{2(\tau_R-\tau_H)}{\theta_{P3}}-\frac{2(\tau_G-\tau_R)}{\theta_R}}e^{-\frac{6}{\theta_R}t_1-\frac{2}{\theta_R}t_2}, k=2,3,4\\
f(g_{go0},t_1,t_2|M_7) &= \gamma_G(1-\gamma_I)\gamma_O\frac{4}{\theta_R\theta_{P2}}e^{-\frac{2}{\theta_{P2}}t_1-\frac{2}{\theta_R}t_2}, t_1<\tau_{P1P2}-\tau_H\\
f(g_{go1},t_1,t_2|M_7) &= \gamma_G(1-\gamma_I)\gamma_O\frac{4}{\theta_R\theta_{P1P2}}e^{-\frac{2(\tau_{P1P2}-\tau_H)}{\theta_{P2}}}e^{-\frac{2}{\theta_{P1P2}}t_1-\frac{2}{\theta_R}t_2}, t_1<\tau_R-\tau_{P1P2}\\
f(g_{go2},t_1,t_2|M_7) &= \gamma_G(1-\gamma_I)\gamma_O\frac{4}{{\theta_R}^2}e^{-\frac{2(\tau_{P1P2}-\tau_H)}{\theta_{P2}}-\frac{2(\tau_R-\tau_{P1P2})}{\theta_{P1P2}}}e^{-\frac{2}{\theta_R}t_1-\frac{2}{\theta_R}t_2}, t_1<\tau_G-\tau_R\\
f(g_{gok},t_1,t_2|M_7) &= \gamma_G\gamma_I(1-\gamma_O)\frac{4}{{\theta_R}^2}e^{-\frac{2(\tau_{P1P2}-\tau_H)}{\theta_{P2}}-\frac{2(\tau_R-\tau_{P1P2})}{\theta_{P1P2}}-\frac{2(\tau_G-\tau_R)}{\theta_R}}e^{-\frac{6}{\theta_R}t_1-\frac{2}{\theta_R}t_2}, k=\ 3,4,5\\
f(g_{gio0},t_1,t_2|M_7) &= \gamma_G\gamma_I\gamma_O\frac{4}{{\theta_R}^2}e^{-\frac{2}{\theta_R}t_1-\frac{2}{\theta_R}t_2}, t_1<\tau_G-\tau_R\\
f(g_{giok},t_1,t_2|M_7) &= \gamma_G\gamma_I\gamma_O\frac{4}{{\theta_R}^2}e^{-\frac{2(\tau_G-\tau_R)}{\theta_R}}e^{-\frac{6}{\theta_R}t_1-\frac{2}{\theta_R}t_2}, k=1,2,3
\end{aligned}
\tag{S1}
$$

Supplementary Figures

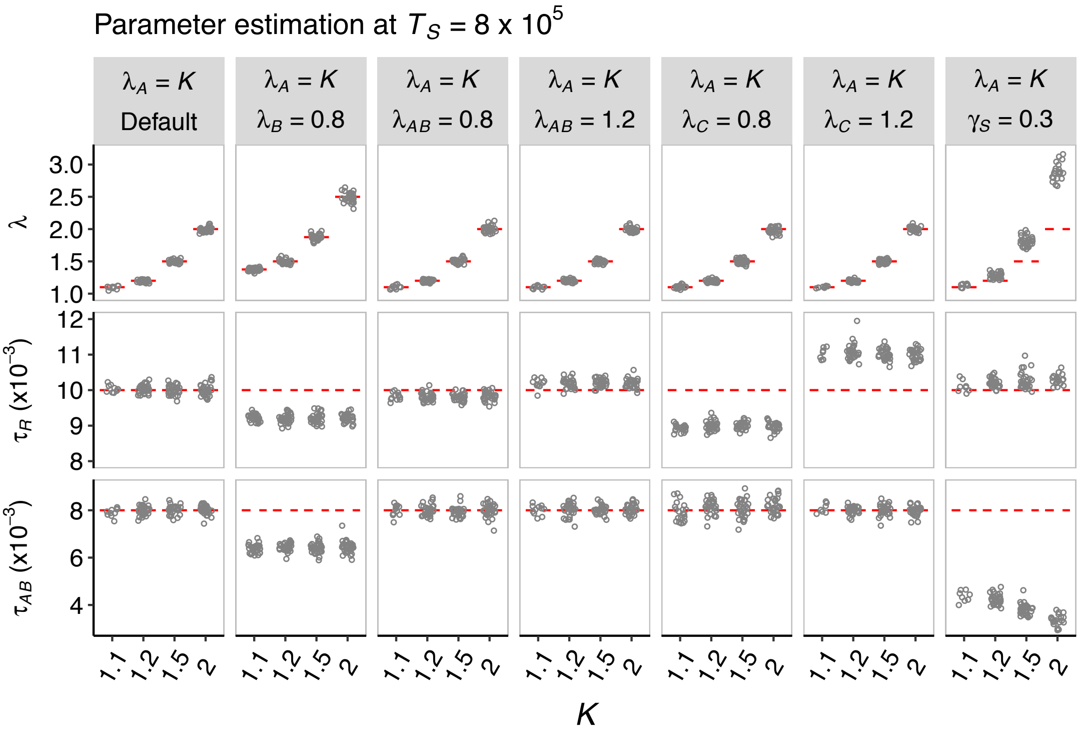


Figure S2. Parameter estimation at $T_{AB} = 8 \times 10^5$ under various rate-variation scenarios in Figure 3a.

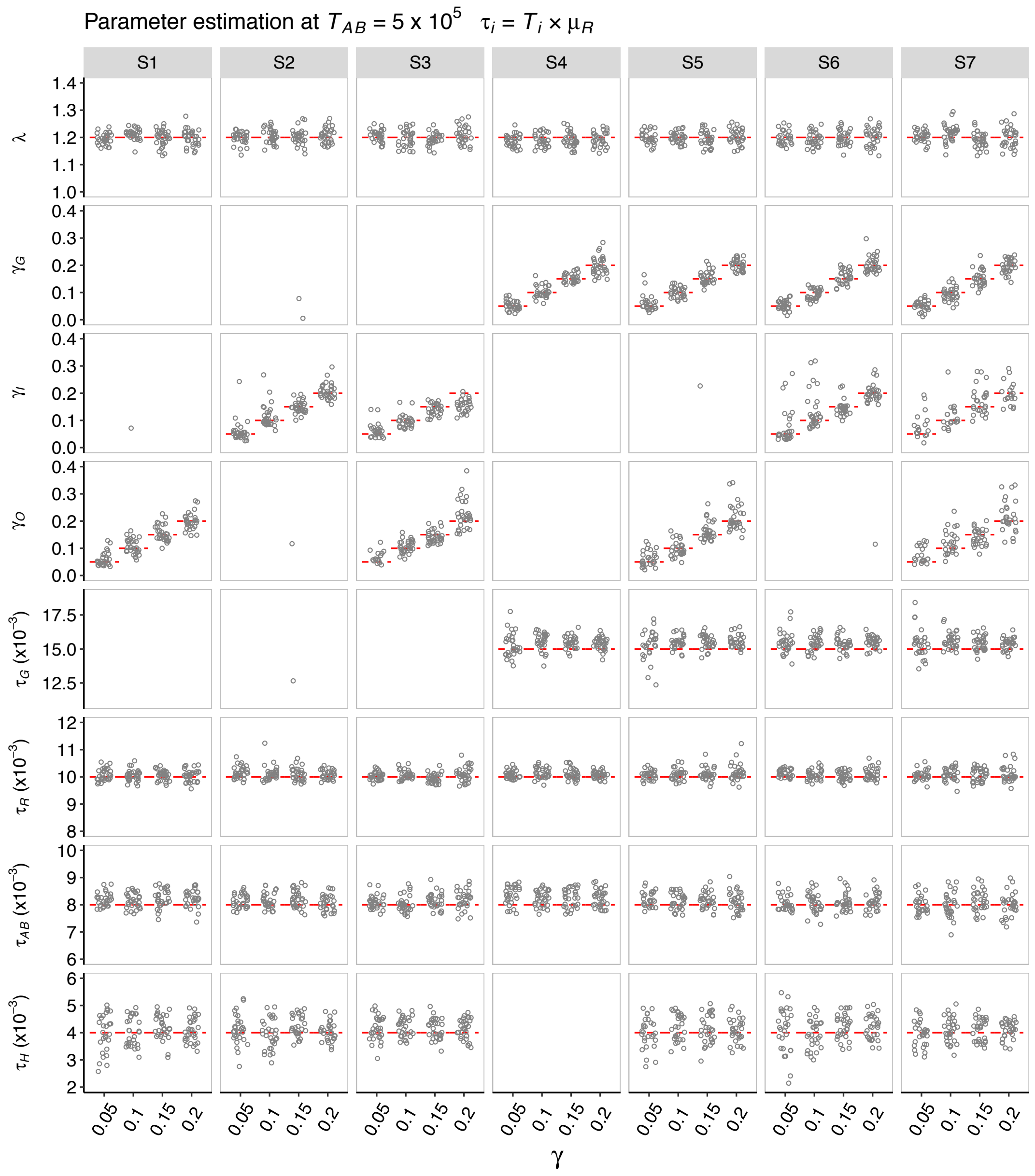


Figure S3. Parameter estimation at $T_{AB}$ = 8 × $10^5$ under various introgression scenarios in Figure 4a.
